# Sharq, A versatile preprocessing and QC pipeline for Single Cell RNA-seq

**DOI:** 10.1101/250811

**Authors:** Tito Candelli, Philip Lijnzaad, Mauro J Muraro, Hindrik Kerstens, Patrick Kemmeren, Alexander van Oudenaarden, Thanasis Margaritis, Frank Holstege

## Abstract

Despite the meteoric rise of single cell RNA-seq, only a few preprocessing pipelines exist that are able to perform all steps from the original `fastq` files to a gene expression table ready for further analysis. Here we present *Sharq*, a versatile preprocessing pipeline designed to work with plate-based 3’-end protocols that include Unique Molecular Identifiers (UMIs). *Sharq* performs stringent step-wise trimming of reads, assigns them to features according to a flexible hierarchical model, and uses the barcode and UMI information to avoid amplification biases and produce gene expression tables. Additionally, *Sharq* provides an extensive plate diagnostics report for quality control and troubleshooting, including that of spatial artefacts. The diagnostics report includes measures of the quality of the individual plate wells as well as a robust assessment which of them contain material from live cells. Collectively, the innovative approaches presented here provide a valuable tool for processing and quality control of single cell RNA-seq data.

## 1 Introduction

Single cell RNA-seq (scRNA-seq) provides an effective way to explore the transcriptome of heterogeneous cell populations. This has proved especially useful in the investigation of highly dynamic processes such as tissue development, cell differentiation, and cancer. Recent advances in the field such as the introduction of multiple cell barcoding strategies and Unique Molecular Identifiers (UMI) have significantly increased throughput and accuracy of single cell sequencing[1-6]. Commonly used scRNA-seq protocols start by FACS sorting into 384 well plates. This guarantees a high degree of control over the cells that will be sequenced, as well as contributing FACS-derived metrics to the available information.

While new experimental protocols are continuously pushing the boundaries of single cell sequencing, processing of the data also faces a number of challenges. Spurious sequencing events must be eliminated and well barcodes have to be demultiplexed while allowing potential mismatches. Low amounts of starting material necessitate a large number of amplification rounds, exacerbating sequence-specific biases, man dating the use of UMIs. Each of the experimental steps may introduce unexpected artefacts (e.g., there may be technical issues in dispensing cells and/or reagents). It is therefore neccesary to monitor a variety of metrics to help recognize the occurrence of such events and pinpoint their cause.

Recently developed tools can deal with some of these issues, but most are available in isolation and are not integrated in a complete workflow. For example, `UMI-tools` [6] can remove amplification artefacts through UMI de-duplication and account for sequencing errors. `Sircel` [7] can effectively recover misread cell barcodes. Fewer pipelines however integrate these steps into a comprehensive workflow [8, 9].

Here we describe *Sharq* **(S**ingle-cell **H**ierarchical **A**ssignment of **R**eads and **Q**uality control), a new preprocessing pipeline for 3’-end scRNA-seq protocols that make use of well plates (96, 384, or larger). *Sharq* is a pipeline that trims and filters out low quality reads, handles both UMIs and cell barcodes, and produces gene expression tables after mapping the reads to a reference genome and hierarchically assigning them to the features annotated on that genome. Finally, *Sharq* produces a comprehensive QC report, providing several plots and helpful metrics to evaluate the quality of the experiment. In particular, it can robustly identify wells where the amplification reaction failed, as well as estimate which cells contained sufficient material relative to an empty well background.

## 2 Approach

*Sharq* is a preprocessing pipeline designed for 3’-end UMI-based protocols such as CEL-seq2 [10] and MARS-seq [11]. The pipeline handles all necessary steps to go from the output of the sequencing platform, to a gene expression table representing RNA abundances for each gene and cell (see figure 1). Additionally, an extensive plate diagnostics report is produced based on the outputs of all intermediate steps.

**Figure 1:**
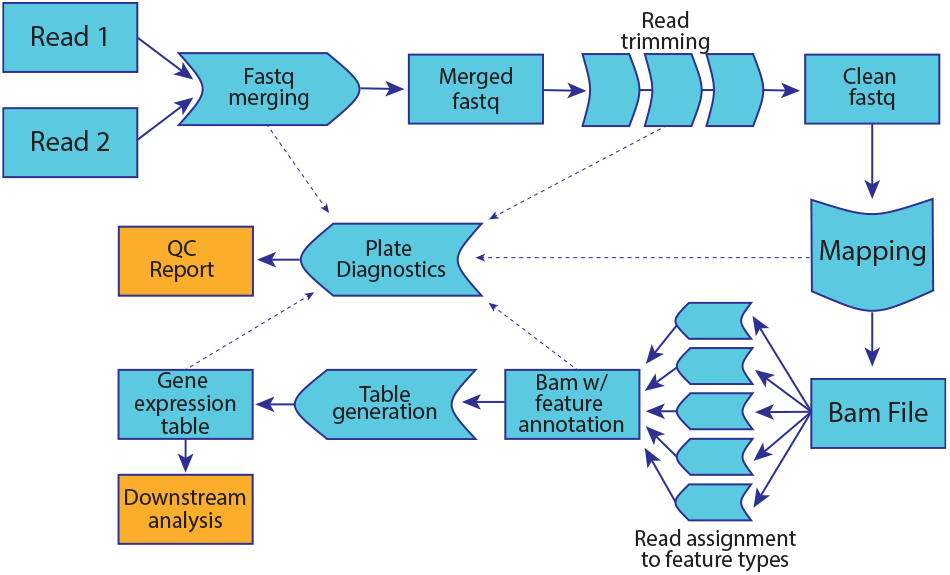
Flowchart representing the steps performed by Sharq. The plate diagnostic tool uses data gathered from intermediate outputs, represented here with a dashed line.

Scripts in *Sharq* are written in Perl, Python 3.6, and R. These scripts are tied together by a pipeline written in Workflow Definition Language (WDL) [12] and can be invoked through a WDL execution engine. This engine supports google cloud solutions and a variety of HPC implementations such as SGE. The entirety of the code and the list of dependencies can be found on bitbucket (https://bitbucket.org/princessmaximacenter/Sharq).

### 2.1 Preprocessing

In Sharq-compatible input files, each read-pair is marked by two distinct barcodes of variable length: a Cell Bar-Code (CBC) that defines the well of origin and a UMI used to detect and correct PCR amplification artefacts. However, while CBC and UMI are stored in read 1 of each pair, the DNA insert of interest is stored in read 2. The first step of the pipeline therefore consists of extracting the barcode information from read 1 and appending it to the header of the corresponding read 2. This preprocessing step produces a single **fastq** file containing both barcode and sequence information.

After UMI and CBC information are integrated, a stringent control quality step is performed on the resulting **fastq** file. A triple trimming approach prevents the inclusion of low complexity and/or low quality reads, either by completely excluding faulty reads, or by trimming away unwanted segments from otherwise high-quality reads (see figure 2). First, each read is probed for the presence of long stretches of A, T, G, or C (18 for A and T, 14 for G and C). The presence of such sequences in the genome is extremely rare, and their inclusion is likely the outcome of spurious sequencing events or other technical artefacts such as inclusion of the poly(A) tail. When these stretches are detected the algorithm removes them along with any nucleotides that follow.

**Figure 2:**
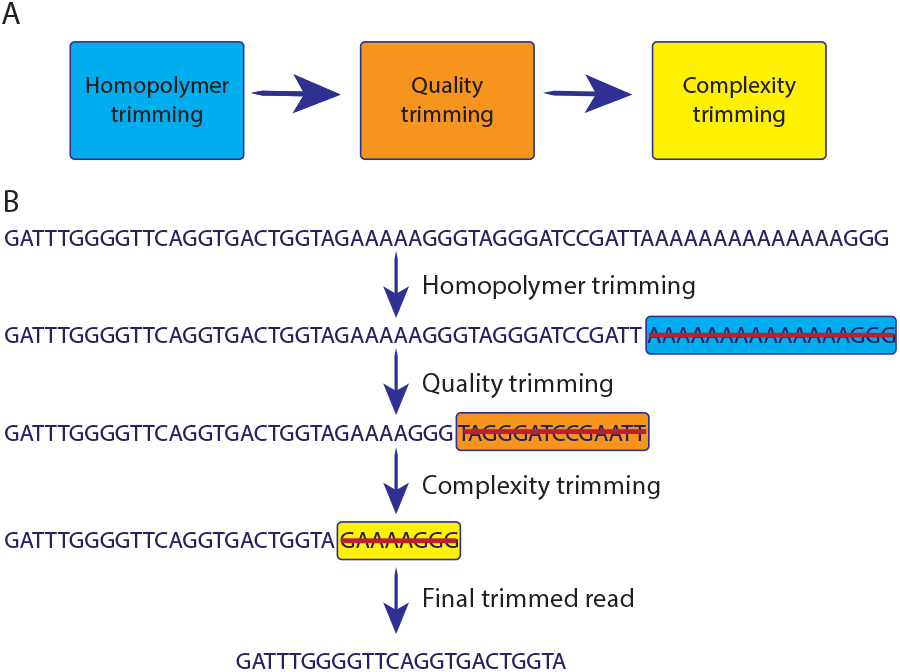
A: Scheme representing the sequential steps that take place during trimming. B: Example of sequential read trimming. First, overly long homopolymer sequences are identified and removed; second, sequencing quality is evaluated and trimmed according to the specified quality threshold; lastly, regions with low complexity are removed.

After the major sequencing artefacts are removed, a second trimming step evaluates the sequence quality of each read in order to further remove low-confidence regions. A common approach is that of trimming all positions below a certain quality threshold from the end of the read, and stopping when a position with sufficiently high quality is encountered. This method, however, often stops prematurely due to the presence of a single high quality position surrounded by low quality ones. To avoid this issue, we use an algorithm originally implemented as part of the short-read aligner BWA [13]. This algorithm makes use of a quality threshold Q to calculate the optimal trimming position according to the following equation:

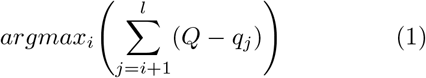

where *i* is a nucleotide position in the read, *Q* is a specified quality threshold, *l* is the length of the read, and *q_j_* represents the quality of the nucleotide at the *j*th position. The position *i* for which this equation is maximal represents the point after which qualities on average start to be lower than *Q*, and can therefore be trimmed away.

The final trimming step targets non-homogeneous low-complexity regions that might have been missed in previous steps. This stage implements a measure of sequence complexity known as CWF [14, 15]:

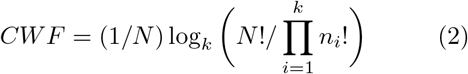

where *N* is the size of the sequence, *n_i_* is the number of nucleotides of the same kind in the sequence, and *k* is the alphabet size (for DNA, *k* = 4). This stage utilises a sliding window approach to calculate a vector of complexities along the whole sequence and then applies the same technique described in (1) to determine if, and how many, nucleotides are to be removed from the read. Default values for window size and complexity threshold are optimized so that protein coding sequences in the human genome are minimally affected by the algorithm, ensuring that reads representing *bona fide* transcripts are preserved.

After trimming, a read is retained only if it still longer than 20 nucleotides.

### 2.2 Mapping and read assignment

After preprocessing and read trimming, passing reads are mapped to a reference genome with the short read aligner STAR [16]. Subsequently, the location of each read must be compared with a reference annotation file in order to determine its feature of origin. Such annotation files sometimes contain two or more overlapping exons that belong to different genes and therefore possess different gene IDs. This makes it difficult for current tools to unambiguously assign the correct feature to reads overlapping these exons, as they are equally likely to have come from either feature. A striking example of this phenomenon is the proto-cadherin gamma locus (see figure 3), where twelve distinct proto-cadherin genes share a common last exon. To avoid discarding reads because of ambiguous feature assignments, we decided to aggregate each cluster of overlapping genes under a unique gene ID chosen among its members. Since functional information about a gene is largely represented in Gene ontology (GO) annotation, we chose the cluster ID based on that of the gene associated with the most GO terms^1^. Scripts and documentation to generate the annotation from e.g. the GENCODE **gtf** files are included in the suite.

**Figure 3:**
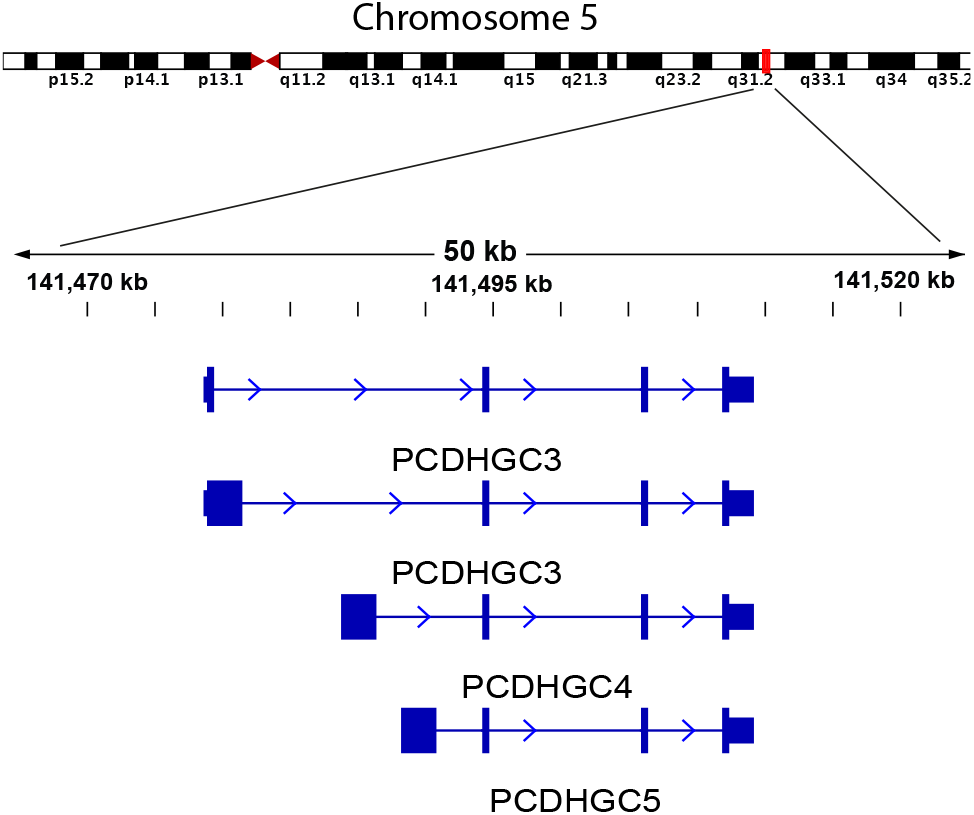
A detail of the protocadherin gamma genomic locus. Four isoforms belonging to three genes are shown. Each isoform consists of four exons (blue boxes), of which three are mutually shared. The complete cluster consists of 22 genes and 50 transcript isoforms, all of which mutually share at least one exon.

Despite these efforts, protein coding genes sometimes still overlap with other feature classes (e.g. long non-coding RNAs). Such genes run the risk of being ignored by the feature assignment algorithm because of their ambiguity. For this reason, we decided to independently assign reads to several classes of features (using `featureCounts` [17]) and later choose the most relevant according to the hierarchical model shown in table 1. This scheme can be adjusted, for instance if additional types are required for features representing pre-mRNA [18, 19].

**Table 1:**
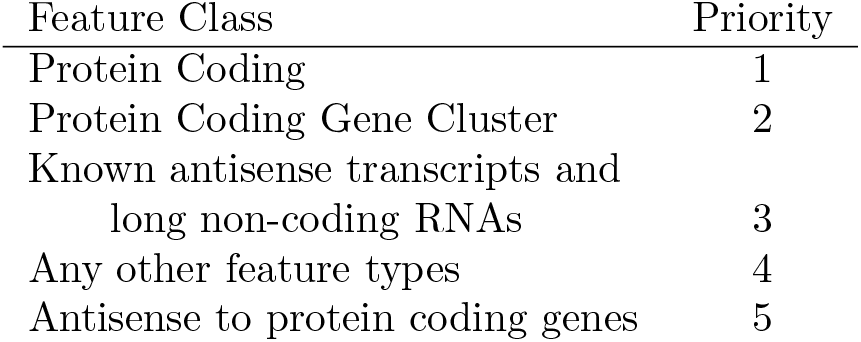
Order of assignment of reads to features. If a read can be assigned to multiple features, only the feature belonging to the class with highest priority will be considered for the purposes of inferring gene expression.

### 2.3 The gene expression table

The final aim of this preprocessing pipeline is a table containing gene expression levels for every individual cell. This requires demultiplexing each read into the corresponding well of origin and ensuring that amplification artefacts are handled correctly.

In *Sharq*, reads are demultiplexed according to provided cell barcodes. However, barcodes are also subject to sequencing errors, which would cause them to differ from the provided ones. To prevent potentially high-quality reads from being discarded we rescue misread barcodes (typically using Hamming distance *≤* 1), but only if they can be unambiguously traced back to a known barcode sequence. E.g., mismatches in certain barcode positions of certain barcodes can be disallowed to make up for shortcomings in the barcode design.

UMI deduplication also takes place at this step, a process that removes amplification artefacts. During the preparation of the library, each recovered transcript is marked by a single UMI sequence. These sequences are usually 6-8 nucleotides long, ensuring that the chance of two distinct transcripts originating from two transcriptional events on the same feature and possessing the same UMI is very low. As a consequence, multiple reads associated with identical UMIs and are assigned to the same feature are likely derived from overamplification of the same original RNA molecule, and are counted only once for the purpose of inferring gene expression. After this step, gene expression tables can be computed using the binomially corrected number of unique UMIs per feature per well as a proxy for RNA abundance [20].

### 2.4 Plate diagnostics

The last step in the pipeline summarises results from all preceding steps in the pipeline and calculates helpful metrics about the plate. A prerequisite for this diagnostic step is the presence of spike-in controls [21] in all wells, as well as the presence of deliberately empty wells. Using these two elements, *Sharq* can accurately indicate: 1) wells where amplification/sequencing reactions failed (dubbed “failed wells”) and 2) wells where the amount of material is significantly above the background given by empty wells (“live wells”).

Failed wells represent technical artefacts where one or more steps leading up to sequencing did not work correctly, ultimately resulting in lack of sequencing reads even for spike-in controls. Failed wells are determined based on the abundance distribution of spikein controls and are discarded from any further consideration. Elimination of failed reactions improves the quality of downstream analyses but it does not guarantee that remaining wells contain biologically meaningful information. The quality of material extracted from single cells is susceptible to a number of issues such as sorting of dead or dying cells which can lead to rapid RNA degradation. We therefore use the abundance of *biological* (i.e. non spike-in) transcripts in *empty* wells as a measure of background noise. This allows us to set an adaptive and robust abundance threshold that identifies wells where biological material is sufficiently above background (see figure 4). Failed and live wells as well as other metrics are overlaid on a schematic representation of the physical plate, allowing identification of potential spatial artefacts (see figure 5). For details of the procedures that determine “failedness” and “liveness” of wells we refer to the functions `goodwells` and `livecellcutoff` of the R package `SCutils` which is also in the bit-bucket repository.

**Figure 4:**
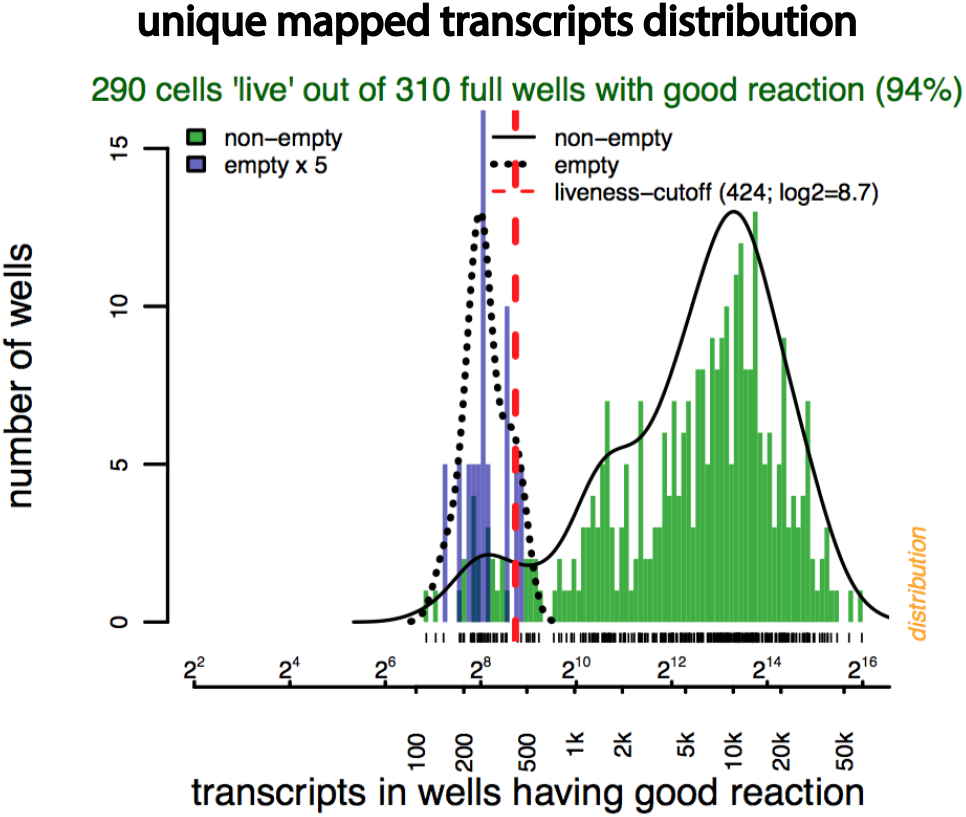
An overview of the distribution of transcript counts, one of the panels of the plate diagnostics report. Distributions are shown both as histograms and as densities, and separately for empty and non-empty wells (the scale of the former is inflated). Wells with transcript counts exceeding the liveness cutoff (red dashed line) are deemed to contain material from live cells.

**Figure 5:**
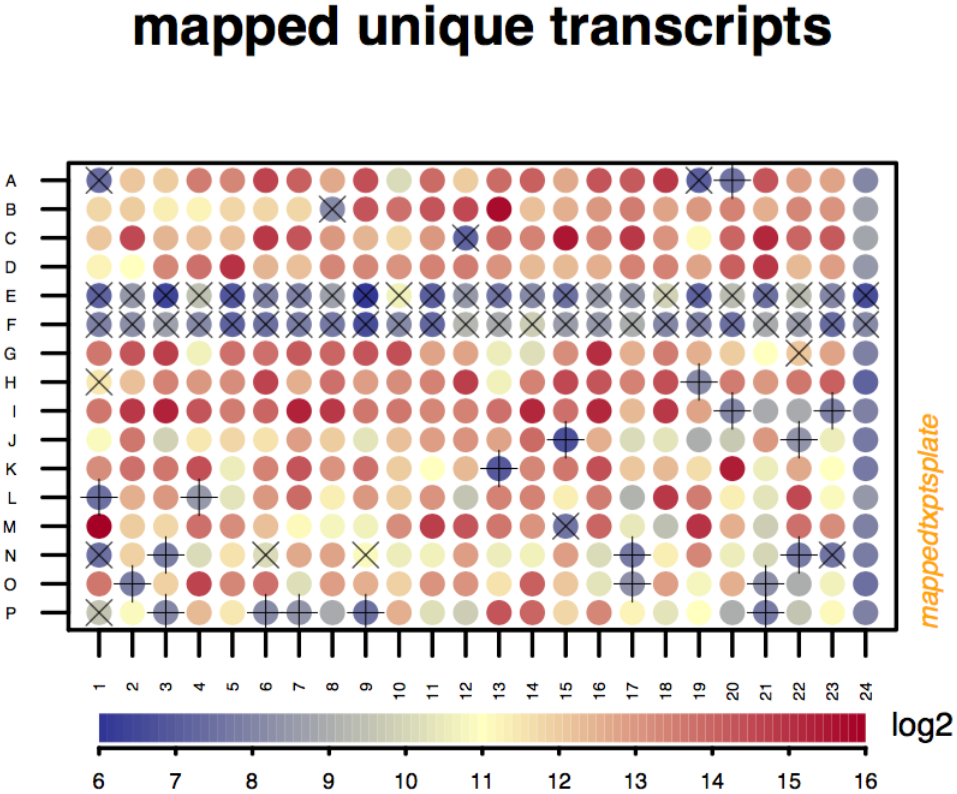
The number of mapped unique transcripts as found in the spatial layout of the plate. Wells where reactions failed are indicated with ×; wells with good reactions but no material from live cells are indicated with +. Column 24 are the wells deliberately left empty; rows E and F had a technical problem. The logarithmic color scale is shown below.

To get an impression of whether deeper sequencing of the same library material is likely to uncover additional genes or transcripts, we go through the **bam** file sequentially and tally the cumulative number of unique genes and transcripts accumulated. The resulting accumulation plots (see figure 6) include estimates of the likely plateau levels, and of the number of reads required to reach 90% of that level. This provides an accurate estimate of the amount of complexity to be gained by additional sequencing on the same sample. Lastly, the plate diagnostics report gives an overview of mapping statistics per feature type and per set of wells, together with: the percentage of transcripts mapping to the mitochondrial chromosome; the distribution of the complexity, of the reads per transcript, and of the number of transcripts per gene; a summary of the most abundant genes and of the most abundant contamination from within and from outside the plate.

**Figure 6:**
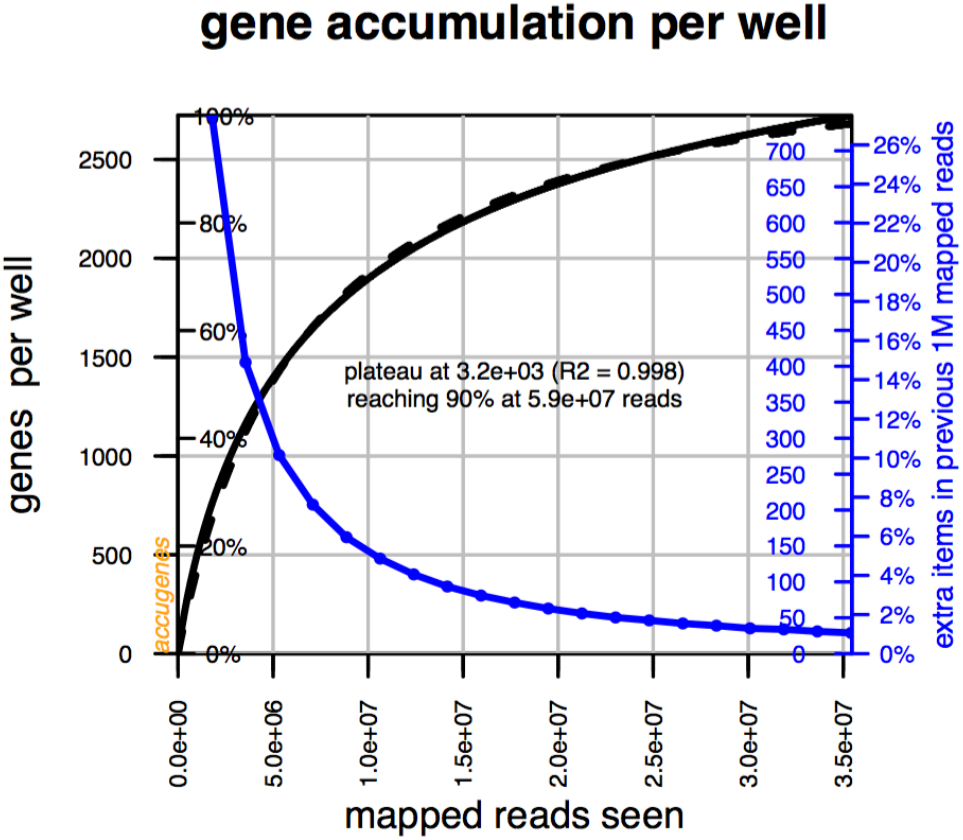
An example of an accumulation plot. The black curve (scale on the left axis) shows how the total number of distinct genes per well increases as more mapped reads are processed while traversing the bam file. The dashed black curve represents the modeled behaviour, yielding the prediction shown in the center. The blue line (scale on the right axis) is the differential of the black line, showing that an additional million reads would probably uncover fewer than 25 extra genes per well.

An example of a full diagnostic report is available as a supplementary document. Detailed documentation on interpretion of the QC rapport is available in the Bitbucket repository.

## 3 Discussion

In this manuscript we describe Sharq, a pipeline designed to process results from the sequencing stage (i.e., the `fastq` files) up to and including the creation of tables with transcript counts per cell and per gene. For each of these steps, statistics are presented to help judge the overall quality of the plate. Further analyses such as normalization, clustering and determining differential expression are left to tools such as RaceID [22] and Seurat [23].

At the current stage of development, Sharq only supports methods that rely on UMIs and sorting into well-plates. However we are considering expanding the codebase to make the Sharq amenable to data produced by 10X Genomics technologies, another leading platform for single cell sequencing. Additionally, we intend to modify our plate diagnostic tool to make comparing statistics across many different plates easier.

Owing to its tremendous promise, the field of single-cell RNA sequencing is advancing rapidly. Consensus on what constitutes best practice is growing, but far from complete. In particular robust measures to establish what are the good cells to consider for further analysis are not yet widely used. The suite of scripts and approaches outlined in this manuscript present a step in that direction.

## 4 Acknowledgments

We thank the members of the Holstege and Kemmeren groups for their support and discussions. This work was financially supported by a grant from the European Research Council.

## Footnotes

1 GO terms that are *Inferred by Electronic Annotation* (IEA) are deliberately left out of this calculation.

